# Zebrafish Information Network, the knowledgebase for *Danio rerio* research

**DOI:** 10.1101/2021.09.22.461443

**Authors:** Yvonne M. Bradford, Ceri E. Van Slyke, Amy Singer, Holly Paddock, Anne Eagle, David Fashena, Douglas G. Howe, Ken Frazer, Ryan Martin, Christian Pich, Sridhar Ramachandran, Leyla Ruzicka, Monte Westerfield

**Affiliations:** The Institute of Neuroscience, University of Oregon, Eugene, Oregon 97403-1254, USA

## Abstract

The Zebrafish Information Network (ZFIN, zfin.org) is the central repository for zebrafish genetic and genomic data. ZFIN expertly curates, integrates, and displays zebrafish data including genes, alleles, human disease models, gene expression, phenotype, gene function, orthology, morpholino, CRISPR, TALEN, and antibodies. ZFIN makes zebrafish research data Findable, Accessible, Interoperable, and Reusable (FAIR) through nomenclature, curatorial and annotation activities, web interfaces, and data downloads. ZFIN is a founding member of the Alliance of Genome Resources, providing zebrafish data for integration into the cross species platform as well as contributing to model organism data harmonization efforts.

## Introduction

The Zebrafish Information Network (ZFIN, zfin.org) is the knowledgebase for the model organism, *Danio rerio* (zebrafish). ZFIN curates, collects, and makes available zebrafish data including genes, gene function, sequences, alleles, mutant and transgenic lines, human disease models, gene expression, phenotype, orthology, sequence targeting reagents, and antibodies. ZFIN is the nomenclature authority for zebrafish genes and alleles and supports the zebrafish research community by providing wiki resources to view antibody and protocol information, as well as ZFIN pages for researchers, laboratories, and companies. ZFIN aims to make zebrafish research data Findable, Accessible, Interoperable, and Reusable (FAIR, Wilkinson et al 2019) by contributing to data annotation standards, the use of biomedical ontologies for data annotations, and persistent identifiers for annotations and metadata. In addition, data are made available at ZFIN through data specific report pages, search interfaces, and download files, as well as through the Alliance of Genome Resources (Alliance, alliancegenome.org). Here we provide an overview of the data types at ZFIN, ways to access data via search interfaces and download files, and data availability at the Alliance.

## ZFIN Data

Genes are at the heart of ZFIN data services, integration, and displays. One of the core services ZFIN provides is gene nomenclature support for the zebrafish community, as well as coordinating nomenclature with other model organism and human nomenclature committees. The gene page serves as the hub of information for genes, providing nomenclature history as well as integrating and providing information about gene expression, mutant phenotypes, associated diseases, gene functions, alleles, and orthologies. Here we describe ZFIN’s gene nomenclature process, the gene page, and data available therein.

### Gene nomenclature

To facilitate unambiguous scientific communication, the ZFIN Nomenclature Coordinator and Zebrafish Nomenclature Committee (composed of active zebrafish researchers) are responsible for approving unique names and symbols for zebrafish loci and transgenic components used in research. A high priority in this process is to ensure that nomenclature reflects a gene’s orthologous relationship to the mammalian (primarily human) genes, and to that end, we regularly coordinate with the HUGO Gene Nomenclature Committee (HGNC) and the Mouse Gene Nomenclature Committee (MGNC). Ensembl (Howe et al., 2021), NCBI (NCBI Resource Coordinators, 2018), and Panther (Thomas et al., 2003) are among the primary resources used in the process of determining gene nomenclature. Nomenclature and orthology are determined using a combination of phylogeny, conserved synteny, and amino acid alignments. Only rarely are coincident expression and functional complementation considered in determining orthology.

The importance of using official, standardized nomenclature cannot be understated. The use of official nomenclature facilitates the discovery and knowledge integration of gene data, making these data accessible and reusable for the scientific community. The proliferation of unofficial, incorrect, or outdated nomenclature leads to confusion, inaccuracy, and difficulties in curating data into and across databases. This can slow research, publication, and curatorial processes, is antithetical to FAIR principles, and can lead to less exposure and fewer citations for researchers. It is critical that authors verify that they are using official gene nomenclature and symbols, as well as obtaining approval for novel gene nomenclature prior to publication. All gene nomenclature can be verified either by searching the ZFIN database or contacting the ZFIN Nomenclature Coordinator.

### Gene page

The Gene page organizes and structures data on the basis of their relationships with a gene, integrating and displaying all data pertinent to the gene so as to facilitate a thorough understanding of gene function (Figure 1). At the top of the gene page, the gene summary section gives basic information about the gene including symbol, links to nomenclature history, name, previously used aliases, a brief computationally generated description of the gene provided by the Alliance (Kishore et al., 2020), and links to gene pages at other databases. The summary section is followed by data sections that present data pertaining to the gene. A navigation pane on the left side of the page allows quick access to the data sections of interest. Citations for the curated data are located at the bottom of the page. Currently, ZFIN contains data on 37,466 genes which are categorized as Protein Coding genes, ncRNA genes, or Pseudogenes.

**Figure 1.**
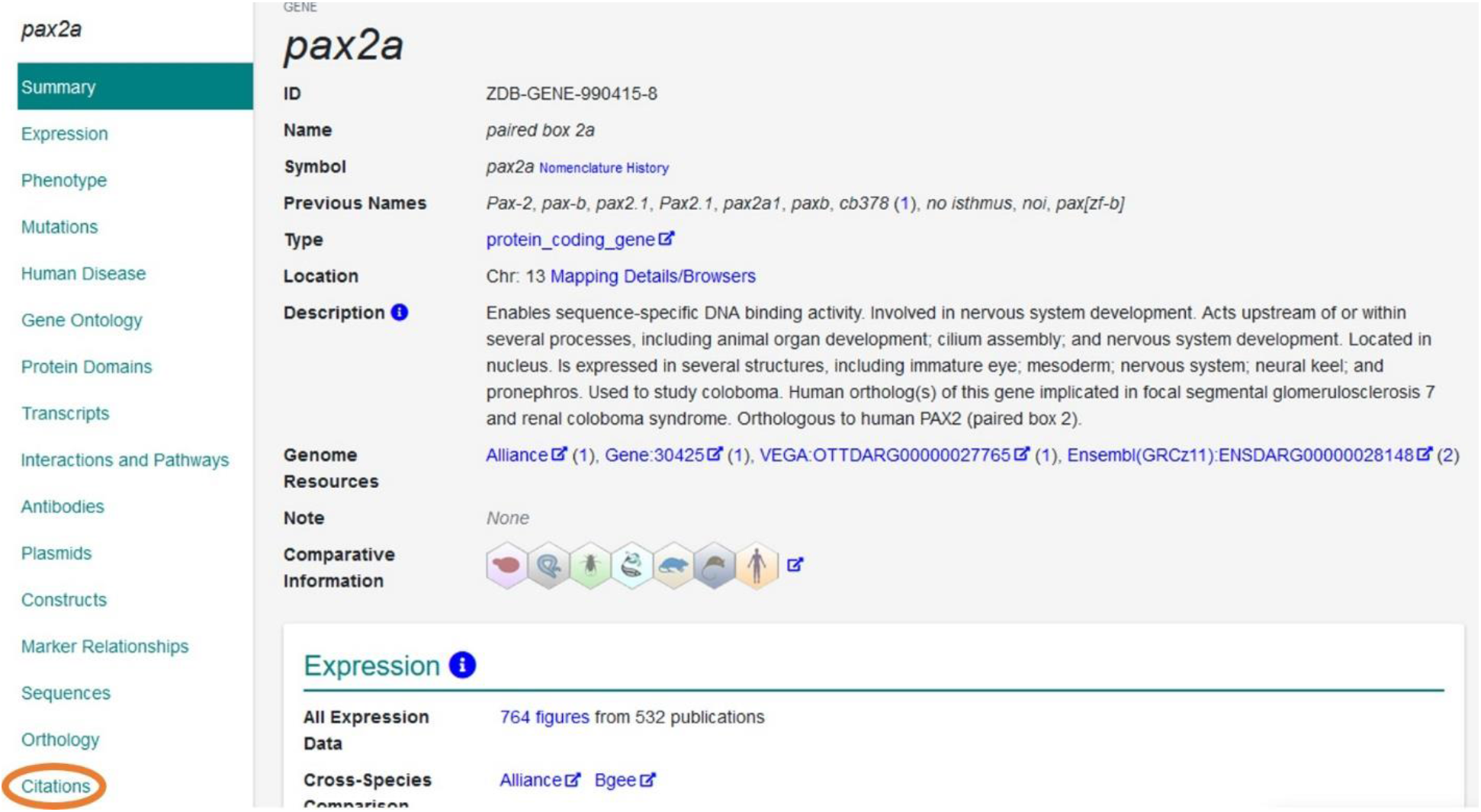
The ZFIN gene page. Web page providing information as it relates to a gene. The top portion provides general information about the gene, and the left navigation panel allows users to navigate directly to areas of interest such as the citation link (circled).

### Gene Expression Section

The ‘Expression’ section on the Gene page focuses primarily on the expression of the gene in a wild-type background (Figure 2). This section begins with an ‘All Expression Data’ link that takes the user to a summary page of all curated gene expression, both in wild-type (WT) and mutant fish, as well as expression under a variety of different experimental conditions. This is followed by links to external sites with cross-species comparison data. The Expression section provides access to high throughput data from GEO and the Expression Atlas, as well as links to data generated in the Thisse large scale WT screen (Thisse et al, 2001). The Wild Type Expression Summary provides a high level visual summary ribbon that denotes the anatomical systems, stages, and GO cellular components that have annotations. Cells in the ribbon that are green indicate the presence of wild-type expression annotations in a system, cellular component, or at a particular stage. Filters above the ribbons can limit expression to *in situs* or to allow the inclusion of expression assays done in reporter lines. To see expression annotation details, the user can click on any cell in the ribbon to see the associated expression annotations (Figure 2). ZFIN curates gene expression using gene symbols, genotypes and strains, the Zebrafish Experimental Conditions Ontology (ZECO, Bradford et al., 2016), the Zebrafish Anatomical Ontology (ZFA, Van Slyke et al., 2014), the Zebrafish Stage ontology (ZFS, Van Slyke et al., 2014), GO cellular compartment ontology (GO-CC, Ashburner et al., 2000, Gene Ontology Consortium 2021), and Spatial ontology (BSPO, Dahdul et al., 2014). Through the use of metadata identifiers and annotation standardization, ZFIN gene expression annotations comply with FAIR standards. ZFIN curates gene expression in both wild-type and mutant backgrounds, as well as in various experimental conditions. The ZFIN gene page displays gene expression only in wild-type backgrounds under standard or control conditions. ZFIN currently has 14,350 genes with expression data and has curated 216,696 zebrafish gene expression assays; of those 134,233 are in wild-type backgrounds with standard or control conditions. In addition, ZFIN has curated 31,964 transgenic reporter assays.

**Figure 2.**
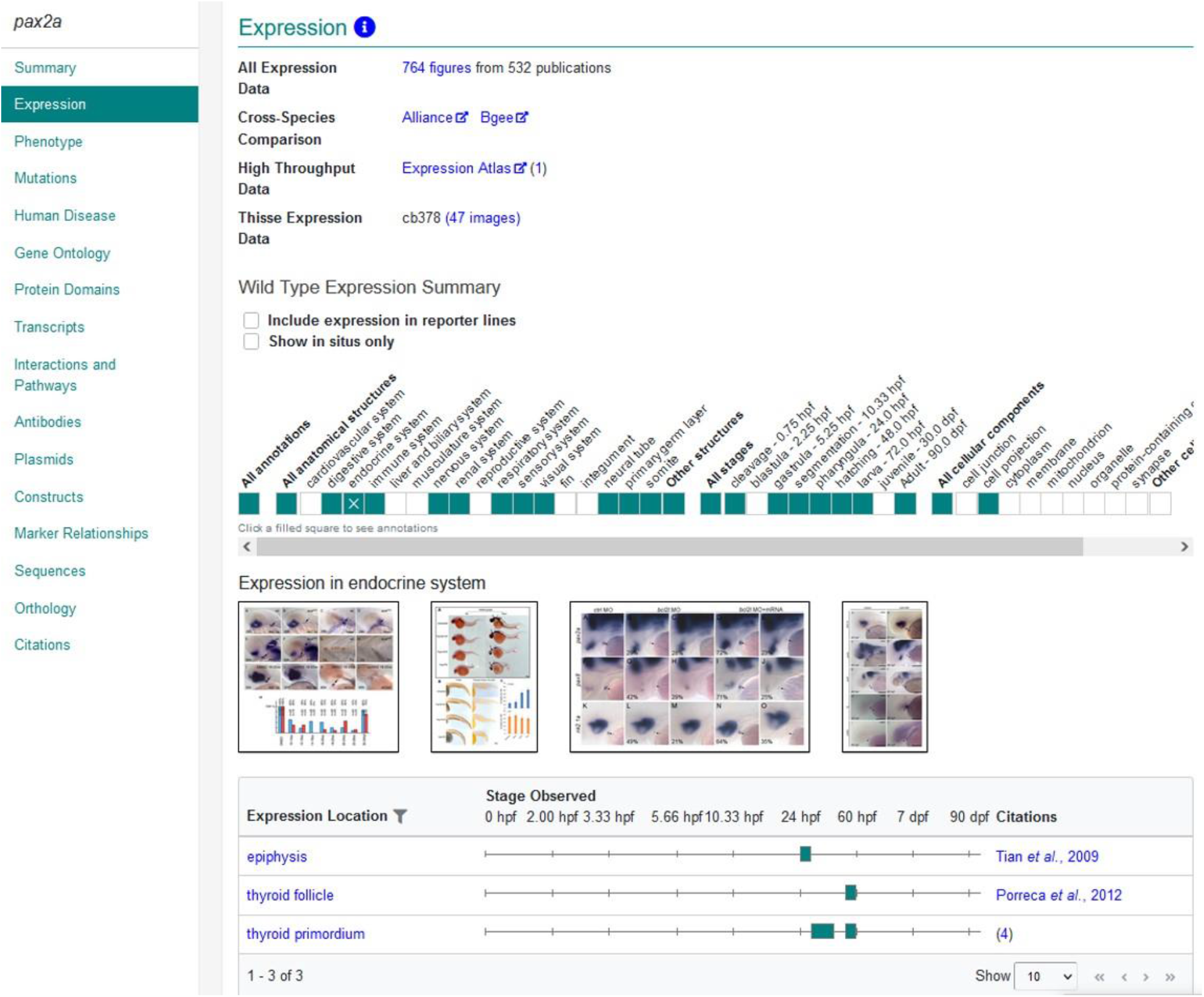
Expression section of the Gene page. Section of the gene page providing information about where and when the gene is expressed. Links to all expression data, Alliance and Bgee for cross-species expression data, high throughput data, and Thisse (Thisse et al, 2001) expression data are provided. The Wild Type Expression Summary provides an overview of expression data, with color blocks indicating where annotations exist. Clicking on shaded boxes opens a table with more detailed gene expression information as shown in the figure.

### Phenotype Section

The ‘Phenotype’ section of the Gene page reports the phenotype of mutant and gene knockdown fish (Figure 3). This section begins with a link to all phenotype data, which when clicked opens a Phenotype Figure Summary page that lists the publications, associated figures, and phenotype annotations in standard or control conditions for the gene of interest. This is followed by a link to the Alliance website for cross-species phenotype comparisons. The Phenotype Summary provides a high level ribbon overview of the anatomical systems, stages, molecular functions, and biological processes that have phenotype annotations. Shaded boxes denote the presence of annotations for the indicated system, stage, biological process, or molecular function. Clicking on the shaded box opens a table view that provides more detailed phenotype annotations (Figure 4).

**Figure 3.**
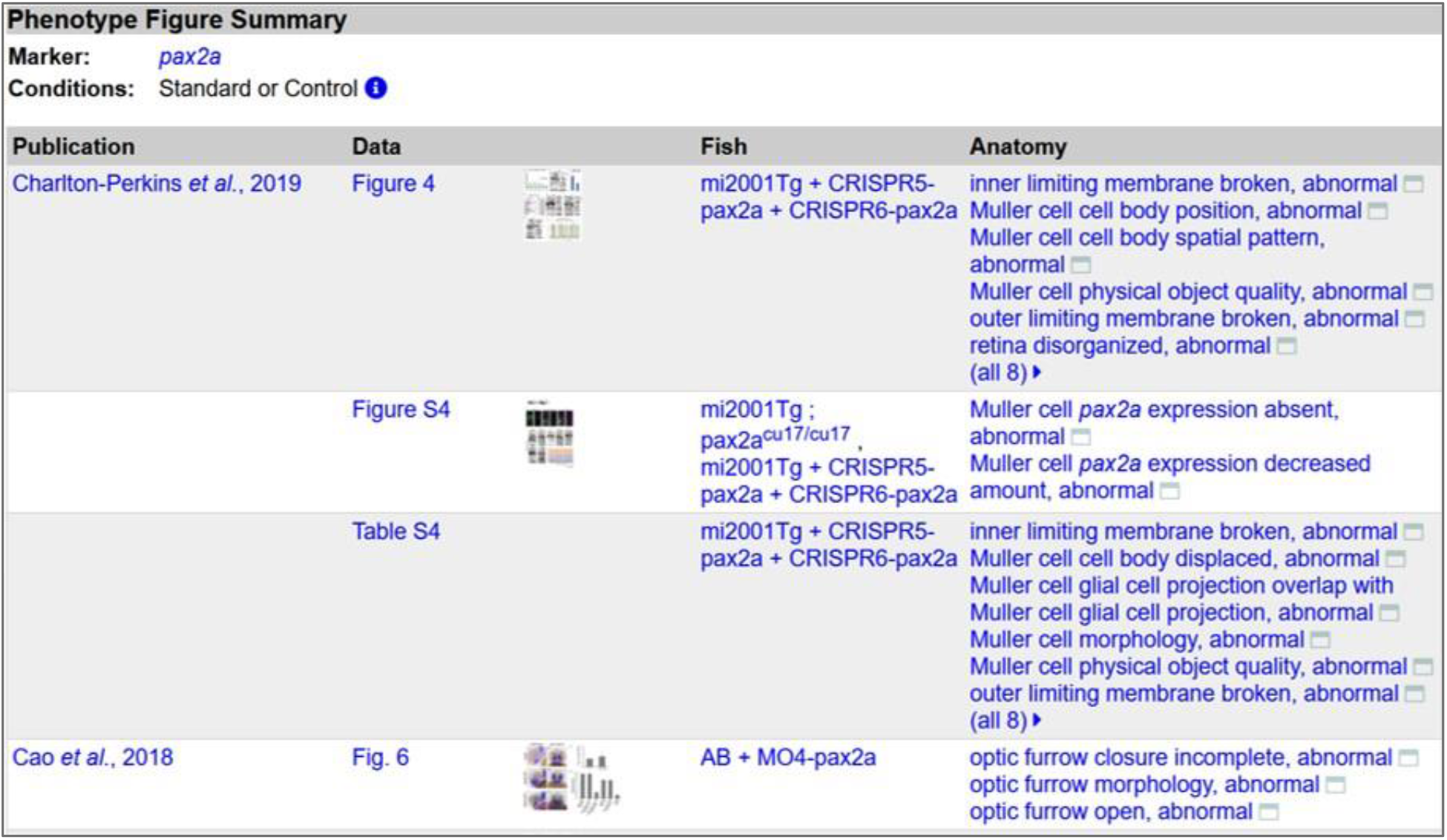
Phenotype Figure Summary. The Phenotype Figure Summary page provides a summary table of, and links to, publications, figures, fish lines, and associated phenotypes for a gene. This page is accessed from the link labeled “All Phenotype Data” on the Gene page.

**Figure 4.**
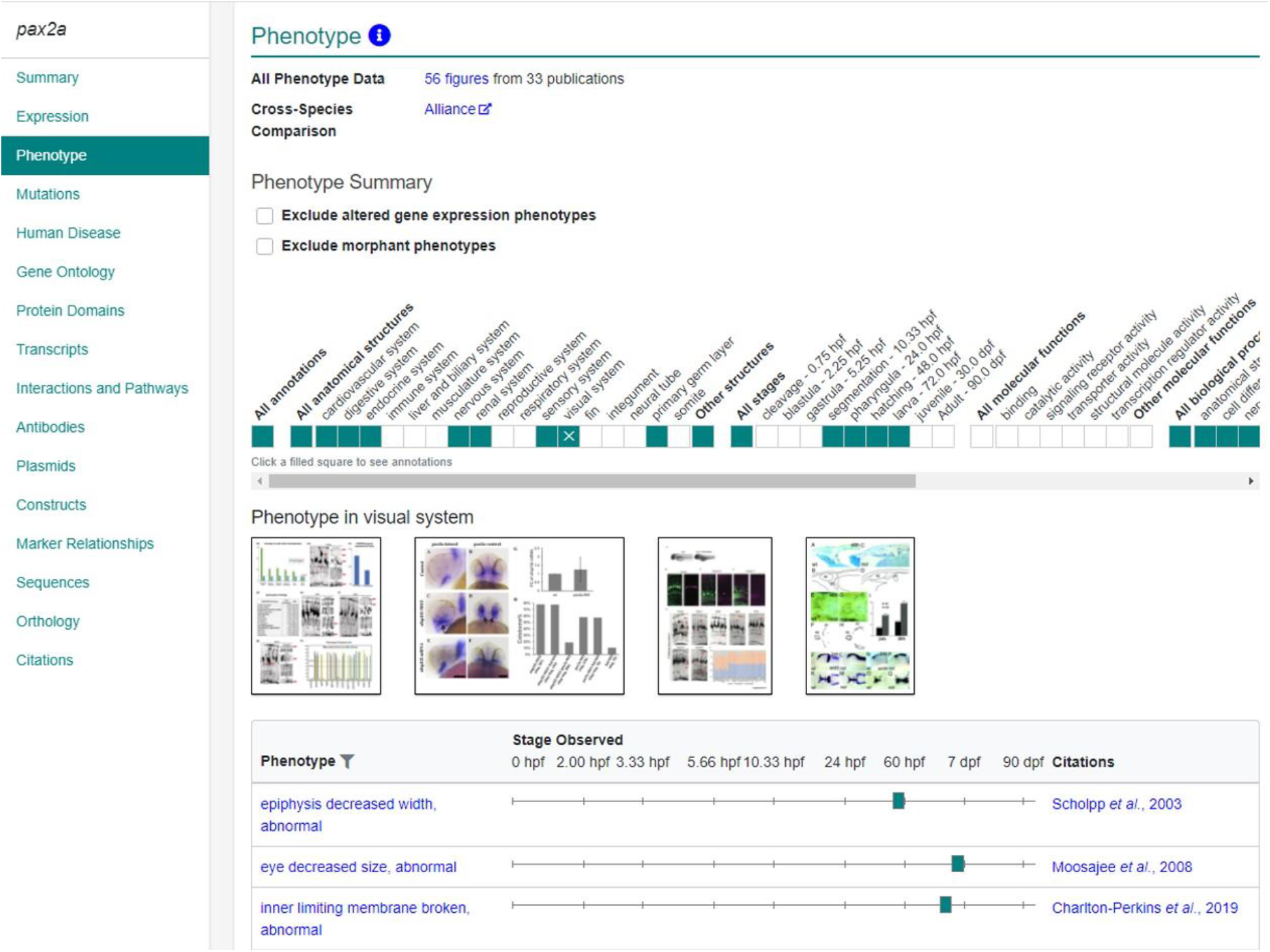
Phenotype section of the gene page. Section of the gene page providing mutant and gene knockdown phenotype information. The Phenotype summary provides a high level overview of systems, stages, molecular functions, and biological processes that have annotations. Clicking shaded boxes provides a table that lists more detailed phenotype annotation information, as shown above.

ZFIN annotates phenotype data using the Phenotype and Trait Ontology (PATO, Gkoutos et al, 2005), ZFA, GO, and Chemical Entities of Biological Interest Ontology (ChEBI, Hastings et al., 2016). A phenotype annotation is composed of the mutant or knockdown fish, the experimental conditions, the stage at which the phenotype is observed, the phenotype statement, and the associated publication. Phenotype statements are composed of entities and qualities assembled in the E+Q syntax (Washington et al., 2009) using the aforementioned ontologies. Phenotype data shown in the phenotype section are from mutant or knockdown fish in standard or control conditions. Phenotype annotations provide core insights into gene function and are utilized to facilitate the understanding of disease processes and outcomes. To facilitate cross-species phenotype integration and knowledge discovery, it is important that phenotype annotations adhere to FAIR principles. To ensure this, ZFIN participates as a core member of the UPheno initiative, which aims to reconcile logical definitions across several model organism phenotype ontologies (Matentzoglu et al., 2018) and has been heavily involved in developing annotation standards and metadata for phenotype annotation (Mabee et al., 2007; Mungall et al., 2010; Köhler et al., 2013). ZFIN has curated 52,305 phenotype statements for single genes under standard or control conditions. There are 4895 genes represented by 7055 alleles with mutant phenotypes, and an additional 2241 genes with morpholino (MO) induced phenotypes where no mutant phenotype has been reported. There are an additional 245 genes with phenotypes reported only in animals injected with CRISPRs (CRISPants).

### Mutations Section

The ‘Mutations’ section of the gene page has two tables, one for ‘Mutants’ and one for ‘Sequence Targeting Reagents’ (Figure 5). This section is intended to provide a high level summary of the alleles and knockdown reagents that have been used to investigate function of the gene. The ‘Mutants’ table summarizes information for the alleles of a gene and includes the allele symbol, which links to the allele page (Genomic Feature Page), the type of mutation, where the mutation occurs within the gene if reported, the transcript consequences, the mutagen, and suppliers if available. ZFIN is the nomenclature authority for mutant lines and records data for zebrafish lines generated by various mutagenesis protocols, including but not limited to transgenic insertions and sequence targeting reagents (STR) such as CRISPRs, TALENs, and morpholinos. To assist in reproducibility, it is important that the nomenclature of alleles and STR knockdown reagents is accurate and distinct prior to publication. This is also critical for line designations/allele numbers. To avoid propagating erroneous transgenic construct nomenclature or duplicate line designations/alleles, authors should search the database or contact the ZFIN Nomenclature Coordinator prior to publication. Each allele record has a line number composed of a one to three letter institution code followed by a number forming a stable identifier for the allele. Institution codes can be found on the ‘Line Designations’ Page (https://zfin.org/action/feature/line-designations). To date, ZFIN has made allele associations for 18890 genes and has records for 72089 alleles, of which 42385 alleles have a one to one relationship to a gene making these alleles useful for gene function analysis.

**Figure 5.**
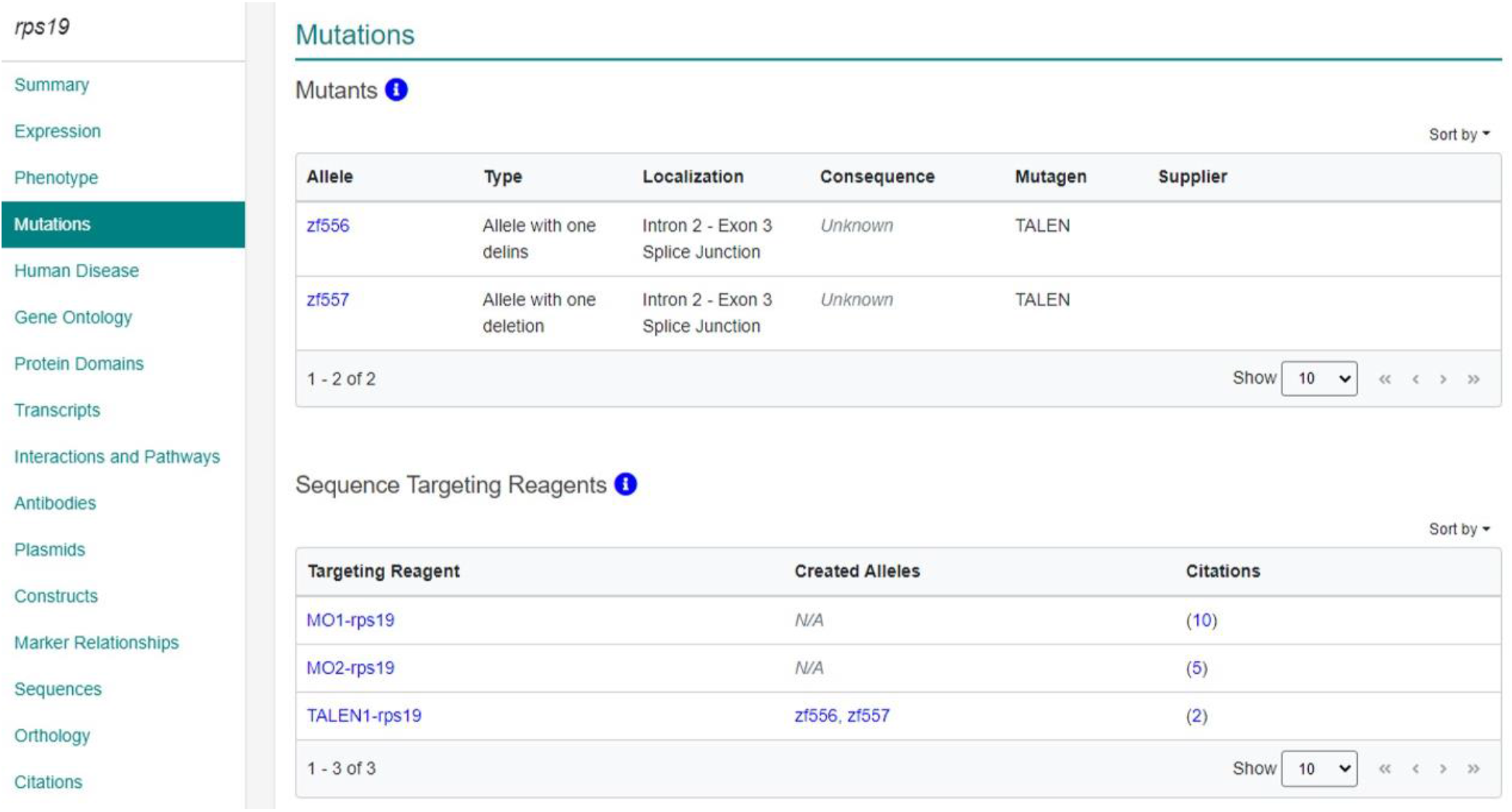
Mutant and Sequence Targeting Reagents section of the gene page. Section of the gene page providing summary information about mutant alleles and STRs for the gene. More detailed information for individual alleles and STRs can be obtained by clicking on the links in the tables.

The ‘Sequence Targeting Reagents’ (STR) table in the Mutations section lists curated MOs, CRISPRs, TALENs used to study a gene (Fig 5). The table lists the STR symbol, which links to an STR specific data page, associated alleles, with links to the Genomic Feature page, and links to references where the reagent is reported. ZFIN facilitates FAIR standards for STR reagents by obtaining STR sequence information from publications, providing distinct nomenclature and identifiers, and reporting this information via user interfaces and download files. In addition, STR sequence targets are verified, and if errors are found authors are contacted to make corrections. Currently ZFIN has records for 10,763 Morpholinos, 6872 CRISPRs and 813 TALENs.

### Human Disease Section

The ‘Human Disease’ section on the Gene page provides two types of disease associations, diseases associated to zebrafish genes via orthology to human disease-causing genes and experimentally verified disease models (Fig 6). The table of diseases associated with the human ortholog lists the OMIM disease name as well as links to the Alliance Disease page, OMIM page, and the corresponding Disease Ontology term (DO, Schriml et al., 2018), with links to the ZFIN DO term page where more information can be found about the disease. Information in this table is produced through computational mappings of ZFIN curated orthologs to human genes and their disease associations from the genemap and mim2gene files from OMIM (https://omim.org/downloads/, Amberger et al 2015). This zebrafish gene to human gene/disease mapping is used to make associations between the ZFIN gene and the DO terms via DO term to OMIM disease mappings in the DO file (Bradford et al., 2017). The Experimental models table provides curated experimental models of human disease. These are fish with a perturbation of the gene of interest that have been experimentally verified to model some or all aspects of a human disease (Bradford et al., 2017) This table contains links to the ZFIN DO term page, the fish line used in the model, the experimental conditions necessary to induce the disease, and a link to the publication where the model was reported. There are additional types of curated disease models available from the DO term pages and the Fish pages. In these models there are either multiple affected genes necessary to model the disease or diseases are modeled via chemical perturbations. Because these models cannot be directly associated with a single gene, they are not included on the individual gene pages. ZFIN currently has 1900 curated disease models and 3606 diseases associated with 5154 zebrafish genes via orthology with 4078 human genes. Individual DO term pages can be found via the search at the top of any ZFIN page.

**Figure 6.**
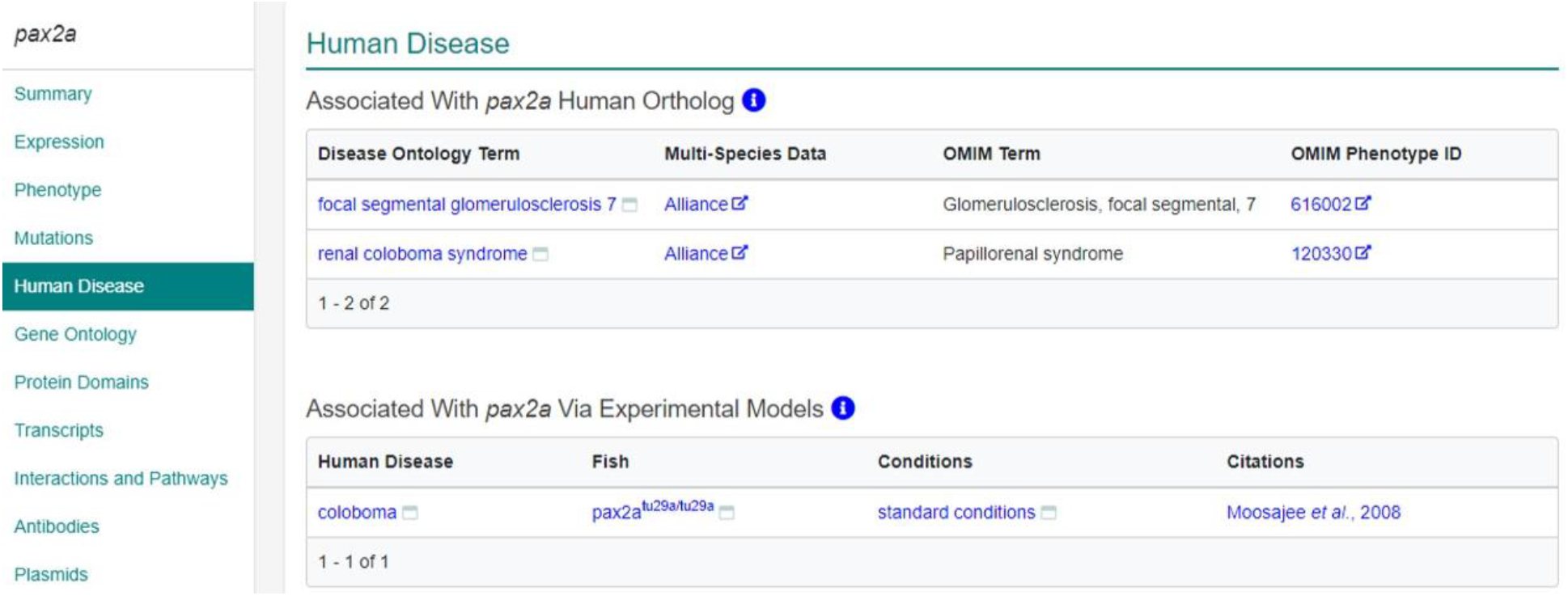
Human Disease section of the Gene page. Provides information about human diseases associated with the gene via orthology to human genes and experimentally validated models of human disease. Links to disease pages provide more detailed information about the disease, associated genes, and experimental models.

### Gene Ontology Section

The ‘Gene Ontology’ section of the gene page reports the roles of a gene or gene product as described by the Gene Ontology (GO) term annotations for the gene (Figure 7). These annotations are created by associating a gene or gene product with GO terms that describe molecular functions (MF) that the gene product enables, the biological process (BP) the gene product is involved in, and the cellular component (CC) where the gene product performs its function (The Gene Ontology Consortium, 2019). The GO section has a high level summary ribbon view of the molecular function, biological process and cellular component annotations for a gene. Clicking on shaded boxes opens a table with more detailed GO annotations that include the annotated GO term, evidence code, with/from field, and citation (Figure 7). ZFIN has 23,047 genes with GO annotations.

**Figure 7.**
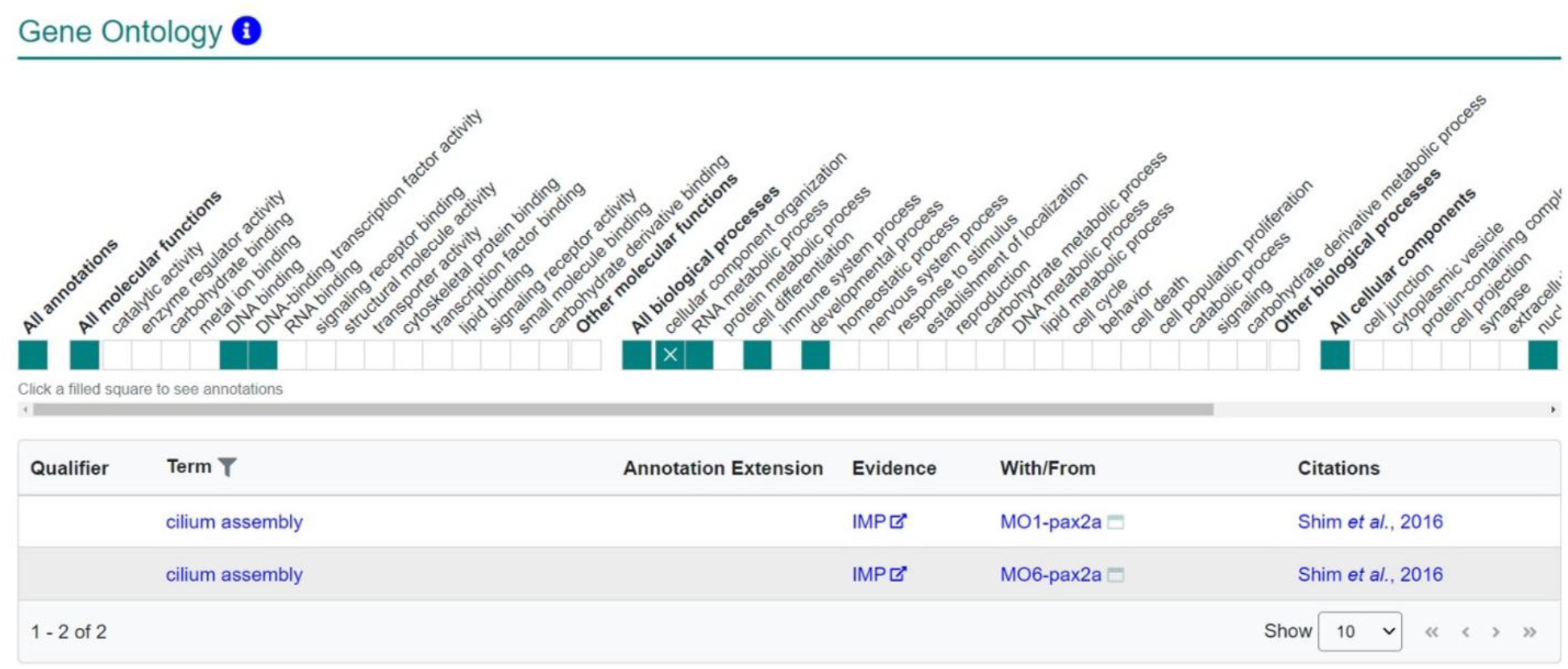
Gene Ontology Section of the gene page. This section provides an overview of the GO molecular function, biological process, and cellular component annotations for a gene. Shaded boxes indicate presence of data; clicking shaded boxes as shown opens a table where more detailed annotation information is provided.

### Orthology and Other Data sections

The ‘Orthology’ section of the gene page provides an overview of the orthology relationships between zebrafish, human, and mouse genes. Orthology displayed in this section has been vetted by an expert curator using conserved synteny, gene family tree analysis, and amino acid alignments. This level of orthology verification is necessary due to the whole genome duplication event in the teleost lineage (Postlethwait et al., 2004) that causes many computational algorithms used for orthology to inadequately identify co-orthologs of mammalian genes. The section links to the orthology section of the zebrafish gene pages at the Alliance where the displays highlight computationally derived orthology and paralogy.

The other sections on the gene page display additional information about the gene and links to related data both at ZFIN and other sites. The Protein Domains section contains links to Interpro domains and lists the UniProt domain details. The transcript section links to several genome browsers as well as displaying the transcript diagrams and links to ZFIN Transcript Pages and withdrawn transcripts. ZFIN links to SignaFish (Csályi et. al 2016) for interactions when known. Antibodies targeting the gene, plasmids with the coding region at Addgene, and constructs containing the gene are linked in their respective sections. There is also a section that links BACs, ESTs, and cDNAs that match the gene as well as a section that links to sequences at GenBank and UniProt.

## Data Accessibility

As the knowledgebase for the zebrafish model organism research community, one of the main goals for ZFIN is to make the data that ZFIN annotates and integrates as accessible as possible. As discussed earlier, ZFIN makes integrated data available on the Gene Page. In addition, data at ZFIN can be searched using the single box search as well as dedicated search forms for specific data types. For computational exploration of data, ZFIN maintains a downloads page that provides access to data specific download files. ZFIN also can provide custom download files upon request. Data provided in download files are FAIR compliant denoting the date the file was generated, as well as all identifiers, symbols, names and ontology terms. ZFIN data are also available at the Alliance. The following describes ZFIN data accessibility via search, download files, and the Alliance web site.

### Search

ZFIN has two options to support user searches for data (Figure 8). The first is a single box search available at the top of the ZFIN home page, as well as the top right of all data pages. The second method uses search forms that allow more specific criteria to be set before beginning the search. Search forms are accessible either on the home page or from the ‘Research’ section of the header under the title ‘Search’ (Figure 8). Both the single box search and the search forms utilize a Solr index of ZFIN data.

**Figure 8.**
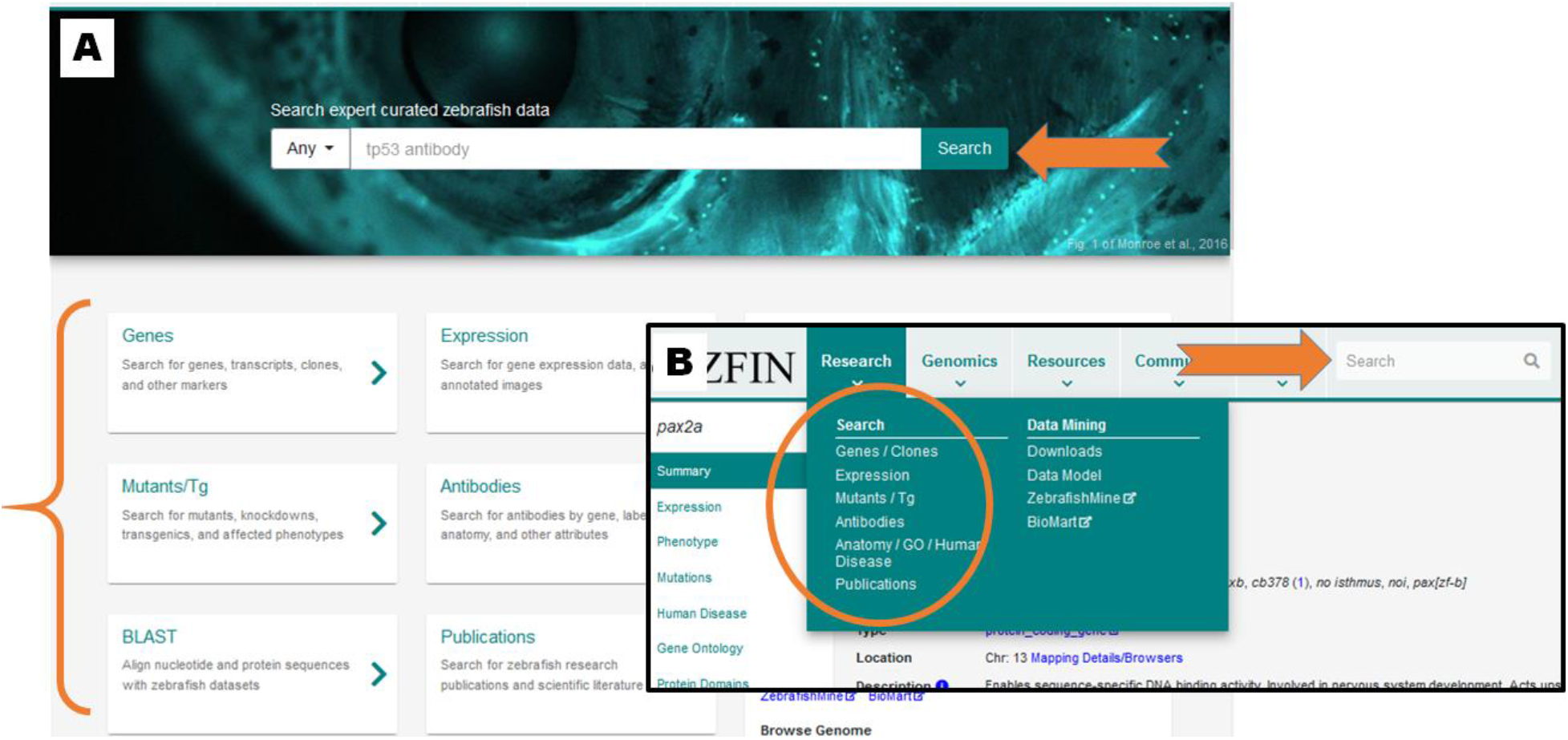
Search ZFIN data. Both Single Box Search and Data Specific Search Forms are available from every page: A) The home page faceted search is accessible from the box centered at the top of the page (left orange arrow) and links to the search forms are in the middle of the page (bracket). B) Search forms are available in the header from the “Research” drop down on every ZFIN page (circled) and the single box search is located on the right side of the header (right orange arrow).

The single box search is a widely-used data exploration paradigm, providing rapid refinement of search results through the use of filters, also called facets (Figure 9). Searches typically begin with text entered into the search box that has an auto-suggest feature to provide a quick lookup of terms, notably genes, alleles, transgenics, and terms from GO, ZFA, and DO, which are extensively used in ZFIN data annotations. Or the search can be initiated with no text which leads to the result pages and allows exploration of all the data at ZFIN.

**Figure 9.**
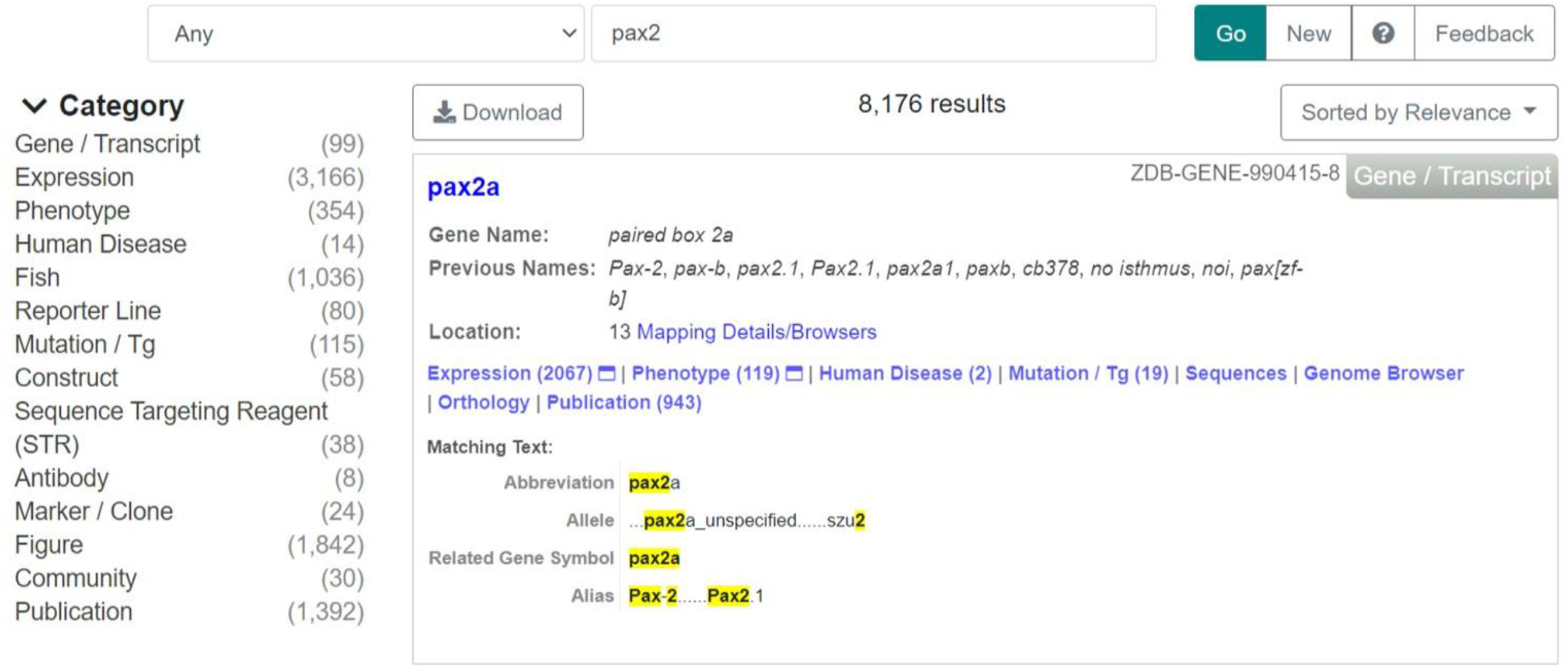
Single Box Search Results page. The results page returned upon executing a query using the single box search. Query results are listed in the main space in the middle, with category facets for filtering results listed in the left panel. The ‘Go’ button executes the search, the ‘New’ button clears the search parameters, the ‘?’ is a help button for single box search questions, and the ‘Feedback’ button sends feedback to the ZFIN Team.

On the result pages data categories are shown on the left side of the search page and can be selected. For any result, stable identifiers found in the results can be downloaded in csv format (Figure 9). Categories include Gene/Transcript, Expression, Phenotype, Fish, Antibody, etc. The number of records in each category that match the entered search string are also shown. Each category has an associated set of attributes (facets). After selecting a category, the left side of the results page is occupied by the “facet panel”. Selecting a facet filters the result to include only data in that facet. Each facet provides a list of values that are valid for filtering the result set. Multiple facets can be selected sequentially. Each of those facets is labelled with a number indicating how many of the current results would remain after selecting that particular facet value. Each time the user selects a facet value, the results and facets are immediately updated. Each of the displayed facet values will be true for at least one of the newly-updated results, ensuring that selecting a facet will never yield zero results.

Single box searching also supports more advanced options including Boolean operators and wild-card searches. For example, searching in the expression category for “fin AND retina” will return expression results that include both fin and retina or their parts. It will not return results that match only fin or only retina or their parts. Asterisk (*) is the wildcard character. At times using a wild-card character can be very helpful to find a larger dataset. For example a search for terms related to the head by using the search string “crani” returns terms with “cranial” or “cranium etc. in them, as expected. Changing the search string to “*crani” returns all those results plus additional terms, which include words like splanchnocranium and chondrocranium that are not found by the original “crani” search.

Search forms have a series of fields pertaining to a topic. ZFIN has search forms for Gene/Clone, Expression, Mutants/Transgenics, Antibodies, Anatomy/GO/Human Diseases, and Publications. These search forms are useful for users looking for a particular subset of data, allowing results to be narrowed down in a single step. The drawback of the search form is that it is possible to eliminate all results. To help ameliorate this drawback, the ZFIN search form remains at the bottom of the search results. Those results can be refined by editing the search form to add more constraints and searching again, or the search can be broadened by removing constraints.

### Download files

ZFIN makes data available for programmatic consumption via download files at https://zfin.org/downloads. These files are accessible from the “Downloads” link found in the “Research” section of the header and from the “Downloads” link in the footer of every page. Downloadable files are organized by categories that represent the data that can be found within the associated files. Clicking on the “Column Headers” shows what is in the file, and clicking on the file name opens the entire file. The file format is listed under “Download with Header” and clicking on that column allows the file to be saved to your computer. An archive of download files is available at https://zfin.org/downloads/archive. The files have been archived since March 28, 2012.

For users who are interested in expression and phenotype data, the most useful download files are “Expression Data for Wild-type Fish” and “Phenotype of Zebrafish Genes”, respectively. The file “Expression Data for Wild-type Fish” is the compilation of all gene expression data for individual genes in WT fish under standard conditions. The “Phenotype of Zebrafish Genes” file contains phenotypes that are observed under “standard” or “control” conditions and have only mutation or knockdown of a single gene. ZDB-PUB to PMID and title mappings are located in the “ZFIN Bibliography” file.

There are several things to keep in mind when analyzing expression and phenotype that are curated using ontology terms. The first is that expression or phenotype listed for a structure means only that the expression or phenotype was observed somewhere in the structure. For expression annotations listing “whole organism” as the structure, this means simply that there was expression somewhere in the fish.

The second similar caveat is that when a stage range is provided for expression or phenotype and a structure is named, that expression or phenotype in that structure was observed sometime during that time period. The last thing to be aware of when using complex data files is that when an abnormal phenotype is unexpectedly absent or expected expression is *not* observed, then those data are curated as such and identifiable by a flag in the file.

For other expression and phenotype data, many different download files are needed. The files are keyed on unique identifiers. The unique identifiers are either ZFIN object identifiers that begin with ‘ZDB-’ followed by the data type and a number, or ontology identifiers that have the ontology prefix and a colon followed by a number. Many of the downloads files are equivalent to a single database table so the Data Model, linked from the home page, is useful for understanding how the files should be linked together. The ontologies referred to in the files can be found at http://www.obofoundry.org/

### ZFIN data at the Alliance

ZFIN is a founding member of the Alliance of Genome Resources (alliancegenome.org, Alliance of Genome Resources Consortium, 2019) which has the primary mission of developing and maintaining comparative genome information resources that facilitate the use of multiple model organisms to understand the genetic and genomic basis of human biology and disease. As of the Alliance 4.1 release (August 2021, see Genetics this issue), ZFIN contributes the following data to the Alliance: genome data, wild-type expression, phenotype data, mutant and transgenic alleles, variants, disease models, and orthology. Through the integration of model organism gene data, the Alliance provides comprehensive gene orthology data. The Alliance utilizes ZFIN data, and other model organism database data, to produce species specific genome browsers that display the genome, genes, and variants. In addition, the Alliance uses ZFIN data to produce zebrafish specific pages like gene and allele, and provides links to data pages at ZFIN including the gene page, allele page, fish page, and disease page. As discussed previously, reciprocal links are provided from ZFIN pages to appropriate Alliance data pages. The Alliance makes ZFIN data available along with data from all the participating model organism databases via either swagger APIs from the Alliance (https://www.alliancegenome.org/api/swagger-ui/) or as download files (https://www.alliancegenome.org/downloads).

## ZFIN technical implementation

The ZFIN technical stack is substantially unchanged since our last publication (Howe et al 2021). In brief the ZFIN web architecture is primarily written in Java using the Spring Framework and served by JSP in Apache/Tomcat. The ZFIN user interface is implemented using React, GWT, Angular, jQuery and plain JavaScript. Groovy, SQL, and Perl are used to process and load bulk data. Hibernate serves as the object-relational mapping library from Java to the relational PostgreSQL database. ZFIN uses both Solr and Java/Spring to support our search interfaces. Data from papers are entered via a web-based curation interface primarily written in GWT with a few implementations of AngularJS interfaces. The community wiki is powered by Atlassian Confluence software (http://www.altassian.com/software/confluence/). A detailed and browsable view of the current ZFIN data model can be found at http://zfin.org/schemaSpy/index.html. Our current plan is to move from our servers to cloud servers to provide dynamic response to load and smooth our server costs.

## Funding

National Human Genome Research Institute at the US National Institutes of Health [U41 HG002659 (ZFIN) and U24 HG010859 (Alliance of Genome Resources)].

